# A novel MRI-based finite element modeling method for calculation of myocardial ischemia effect in patients with functional mitral regurgitation

**DOI:** 10.1101/822502

**Authors:** Yue Zhang, Vicky Y Wang, Ashley E Morgan, Jiwon Kim, Liang Ge, Julius M Guccione, Jonathan W Weinsaft, Mark B Ratcliffe

## Abstract

**Background:** Functional Mitral Regurgitation (FMR) affects nearly 3 million patients in the United States. Conventional indices are limited for predicting FMR response to coronary revascularization (REVASC). Uncertainty as to which patients will respond to REVASC alone impedes rational decision-making regarding FMR management. Determination of myocardial material parameters associated with ischemic myocardium will address knowledge gaps regarding the impact of ischemia on regional cardiac muscle.

**Method:** We proposed a novel MRI-based finite element (FE) modeling method to determine the effect of ischemia on myocardial contractility. The method was applied to two patients with multi-vessel coronary disease and FMR and one healthy volunteer. Cardiac MRI (CMR) included cine-MRI, gadolinium-enhanced stress perfusion, late gadolinium enhancement (LGE), and non-invasive tagged MRI (CSPAMM). The left ventricular (LV) FE model was divided into 17 sectors. Sector-specific circumferential and longitudinal end-systolic strain and LV volume from CSPAMM were used in a formal optimization to determine the sector based myocardial contractility, T_max_.

**Results:** The FE optimization successfully converged with good agreement between calculated and experimental end-systolic strain and LV volumes. Specifically, the optimized *Tmax_H_* for Patient 1, Patient 2, and the volunteer was 336.8 kPa, 401.4 kPa, and 259.4 kPa and α for Patient 1 and Patient 2 was 0.44 and 0.

**Conclusion:** We developed a novel computational method able to predict the effect of myocardial ischemia in patients with FMR. This method can be used to predict the effect of ischemia on the regional myocardium and promises to facilitate better understanding of FMR response to REVASC.

## 1 Introduction

Functional mitral regurgitation (FMR) is a leading cause of valvular heart disease. FMR occurs in nearly 3 million people in the United States, more than 400,000 of whom have advanced (≥ moderate) MR (Gorman et al., 2003; Groarke et al., 2017). Left ventricular ischemia is a potential causal factor for FMR, as evidenced by animal studies showing that ischemia induction via coronary ligation yields acute MR which resolves after restoration of coronary patency (Kono et al., 1992; Messas et al., 2001). Although these data support the notion that FMR can be effectively treated when ischemia is relieved, clinical uncertainty persists as to whether ischemia is a primary determinant of FMR response to coronary revascularization in the context of confounding variables such as myocardial infarction (MI) and left ventricular (LV) chamber remodeling. As evidence of this, clinical studies have shown that in approximately half of patients, MR improves with coronary revascularization (REVASC), whereas in the remainder MR persists or worsen (Aklog et al., 2001; Penicka et al., 2009; Kang et al., 2011). Uncertainty as to which patients will respond to revascularization impedes rational decision-making regarding MR management.

Finite element (FE) modeling has the unique potential to discern the impact of ischemia on MR in context of concomitant LV remodeling and infarction. Prior studies by our group (Walker et al., 2005; Wenk et al., 2011) and others (Bogen et al., 1980; Genet et al., 2015) have focused on the impact of MI alone on FMR but without incorporation of ischemia effect into computational models. Other FE modeling studies have incorporated ischemia into simulations focused on arrhythmogenesis (Wang et al., 2013; Mendonca Costa et al., 2018) without incorporation of LV geometry or function. McCulloch described the implementation of the coronary blood flow myocardial function relationship in a FE framework (McCulloch and Mazhari, 2001). However, to date, modeling of ischemia effect on LV contractility as occurs in infarcted and hibernating myocardium (Wijns et al., 1998) has not been described in the context of advanced LV remodeling with multivessel coronary disease (CAD).

New advances in non-invasive imaging enable both LV ischemia and infarction to be assessed with high precision, facilitating inclusion of such data into FE models. Cardiac magnetic resonance (CMR) has been well validated for both LV remodeling and tissue characterization. Cine-CMR has been employed as a reference standard for LV chamber size and function (von Knobelsdorff-Brenkenhoff et al., 2017). Stress perfusion (SP) CMR has been shown to provide high diagnostic accuracy for ischemia as can occur with obstructive CAD (Bruder et al., 2013). Late gadolinium enhancement (LGE) CMR provides near exact agreement with histopathology evidenced myocardial infarction (Kim et al., 1999; Kim et al., 2000) and CMR with non-invasive tags (CSPAMM) has been widely used in assessing LV contractility (Ryf et al., 2002). This study leveraged multiparametric CMR data to develop and validate a novel computational method to predict the effect of myocardial ischemia on regional left ventricular function in patients with FMR.

## 2 Methods

Two patients and one healthy volunteer were prospectively enrolled in a protocol examining FMR-associated remodeling. Imaging was performed at Weill Cornell Medical College (New York, NY) using 3.0 Tesla scanners (General Electric, Waukesha, WI) (Morgan et al., 2018). The Cornell Institutional Review Board approved this study, and written informed consent was obtained at time of enrollment.

### 2.1 Computational modeling pipeline

The proposed method is summarized in **Figure 1**. The CMR images were used to create the LV & right ventricular (RV) surfaces (**Section 2.3**), to calculate the 3D regional strains (**Section 2.6**), and to configure the LV mechanical parameters (**Section 2.7**). Along with the patient-specific LV and RV pressures, the LV mechanics models were constructed using the LV and RV surfaces, rule-based fiber angles, diastolic and systolic myocardial constitutive relationships and LV myocardial material parameters. A formal parameter optimization framework was then constructed based on the LV mechanics models and the *in vivo* sector-specific circumferential and longitudinal end-systolic strain and LV volume.

**Figure 1.**
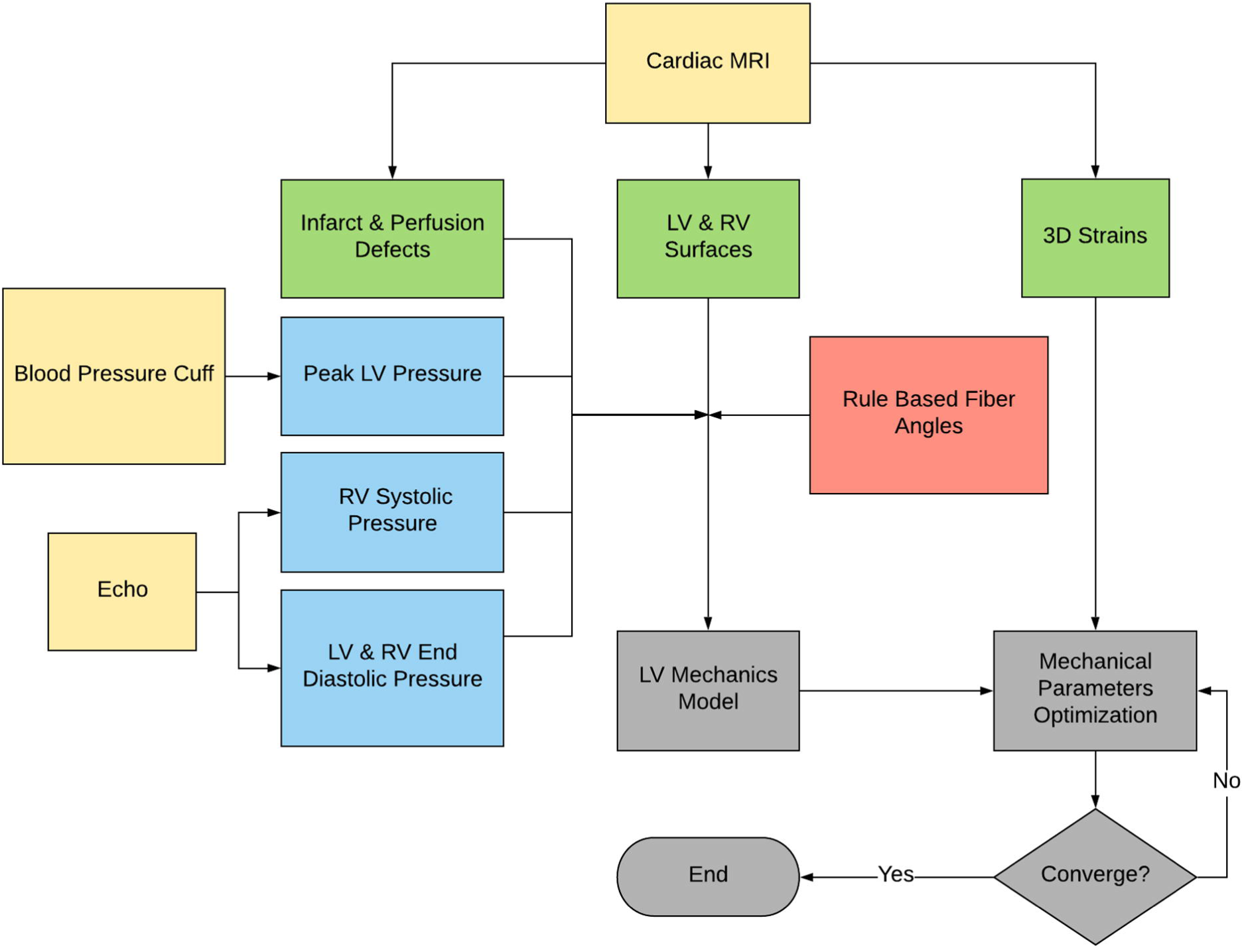
Flowchart of the proposed method. Note that the raw data are in yellow, the derived pressure in blue, the MRI-derived geometry and strains in green, and the FE modeling process in gray.

### 2.2 Image acquisition

Four CMR sequences were collected for each subject. The ventricular structure/function were assessed by Cine CMR with steady-state free precession (**Figure 2A**). The LV ischemia was assessed by the Gadolinium-enhanced first pass Regadenoson-induced SP (4-5 equidistant LV short-axis images) (**Figure 2B**). The LV infarction was assessed by the delayed-enhancement inversion recovery CMR for LGE, 10–30 minutes after administration of gadolinium (0.2 mmol/kg) using a segmented inversion recovery sequence, with inversion time tailored to null viable myocardium (**Figure 2C**). CSPAMM in contiguous LV short and long axis slices (8 mm tag spacing, 10 mm slice thickness, no gap) were performed to measure the myocardial deformation and strain (**Figure 2D**).

**Figure 2.**
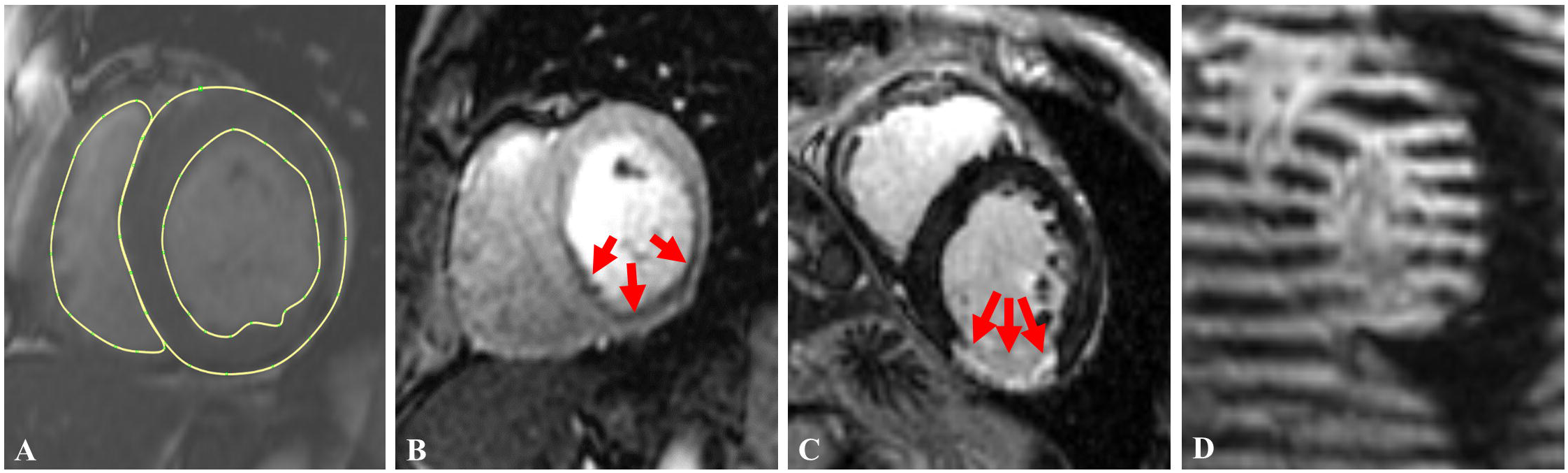
Examples for 4 sequences of images collected in this study: (A) short axis Cine-MRI with LV epicardial and endocardial and RV endocardial contours; (B) a stress perfusion image with arrows indicating the ischemic region; (C) a late gadolinium enhancement (LGE/infarct) image with red arrows indicating the infarcted region, and (D) a short axis tagged image.

### 2.3 Image analysis

#### 2.3.1 LV and RV contouring

The LV (both epicardium and endocardium) short and long axis and the RV (only endocardium) short axis Cine-CMR were contoured at LV early diastolic filling phase (EDF) in a medical image processing platform, MeVisLab (version 2.7.1, Bremen, DE). The EDF phase was defined as the beginning of the opening of the mitral valve. The short axis Cine-CMR were contoured from apex to the mitral valve plane. For this study, only short-axis contours were used for surface generation. The LV and RV contours included the papillary muscle bases but excluded the trabeculae in this study. An example is shown in **Figure 2A**.

In plane shift of the short axis cine-CMR due to breathing resulted in a misalignment of the LV and RV short axis contours. To eliminate this in-plane misalignment, the LV epicardium in three LV long axis cine-CMR (2-, 3- and 4-chamber views) were contoured. A LV epicardial surface was created based on those three long axis contours and this 3D surface had a cross section in each short axis image plane. The centroid of each cross section was calculated and denoted as ***A***. Each LV short axis epicardial contour had a calculated centroid denoted as ***A***′. A transform vector for each short axis contour then was calculated using ***A - A***′. This transform vector for each image plane was applied on both LV and RV short axis epicardial and endocardial contours. After the alignment, the LV epicardial and endocardial and RV endocardial surfaces were created using the aligned short axis contours.

#### 2.3.2 MRI-measured Strains

Regional 3D circumferential and longitudinal strains for each sector were calculated from CSPAMM images using an in-house software as described previously (Morgan et al., 2018). The HARmonic Phase (HARP) method developed by Osman et al. (Osman et al., 2000) was implemented. The implementation details has been reported by Morgan et al. in the Appendix of (Morgan et al., 2018).

### 2.4 LV FE model generation

The LV epicardial and endocardial surfaces were input to TruGrid (XYZ Scientific Applications Inc, Pleasant Hill, California, USA) to mesh the LV as shown in **Figure 3**. Meshes with different density were created. Optimal mesh density was determined as a function of parameter calculation accuracy and calculation time. Custom software (C#, Visual Studio 2017, Microsoft, Redmond, WA) that included a ray casting method (Möller and Trumbore, 1997) to locate the interventricular septum (**Figure 3A**) was used to break the mesh into 17 AHA sectors (Cerqueira et al., 2002) (**Figure 3B**).

**Figure 3.**
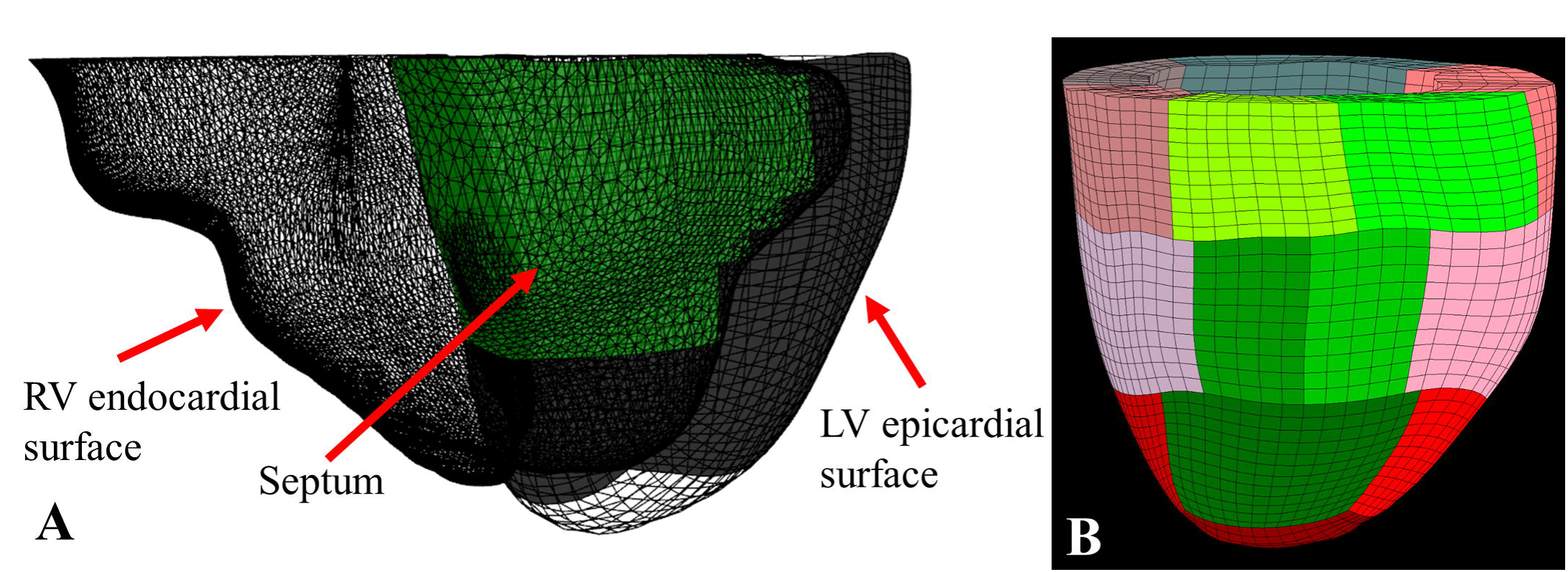
(A) RV endocardial surfaces (triangle mesh), LV epicardial surface (hexahedral mesh), and the septum (in green) determined using a ray-casting method; (B) the final FE model built with 17 sectors with the septal sectors in green.

### 2.5 Loading and boundary condition

The LV systolic pressure was obtained from blood pressure cuff and LV ED and RV ED and ES pressures were estimated from concomitant transthoracic echocardiographic examination data (Ommen et al., 2000). Pressure data is summarized in **Table 1**. Patient-specific LV and RV pressures were applied on the LV endocardial and LV septal epicardial surfaces, respectively. Nodes at the LV top surface were applied with homogeneous Dirichlet boundary conditions. Briefly, basal enpicardial and endocardial nodes were able to slide within the LV valve plane during passive filling and systolic contraction.

**Table 1.** Patient-specific LV and RV pressures.

|  | Patient 1 | Patient 2 | Volunteer |
| --- | --- | --- | --- |
| LV EDP [mmHg] | 20 | 10 | 10 |
| LV ESP [mmHg] | 140 | 140 | 120 |
| RV EDP [mmHg] | 8 | 8 | 8 |
| RV ESP [mmHg] | 28 | 40 | 25 |

### 2.6 Constitutive laws

Passive and active myocardial constitutive laws described by Guccione et al. (Guccione et al., 1991; Guccione et al., 1993) were used in this study. The LV myocardium was modeled with a user-defined material subroutine in the explicit FE solver, LS-DYNA (Liver-more Software Technology Corporation, Livermore, CA). Each FE model was assigned with the rule-based fiber angles with myofiber helix angle varying transmurally from 60° at the endocardium to -60° at the epicardium.

### 2.7 Sector-based myocardial material parameters

The LGE and SP scores were defined as integers from 0 to 4, where 0 represents healthy and 4 the most severe. Passive stiffness, *C*, was determined using the following:

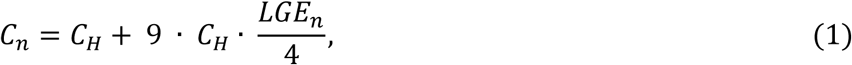

where *C_n_* represents the passive stiffness for each sector and *C_H_* the stiffness of a ‘healthy’ sector with LGE = 0. Note that when LGE = 4 (transmural MI), *C_n_* = 10 *C_H_* which is consistent with our prior work (Wenk et al., 2011).

On the other hand, both LGE and SP were assumed to effect regional contractility, *Tmax*, according to the following linear relationship:

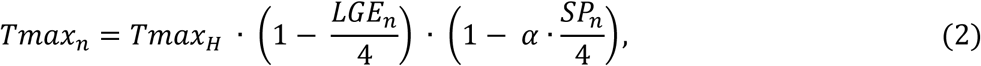

where *Tmax*_*n*_represents the contractility for each sector, *Tmax_H_* the contractility for healthy sector with LGE and SP = 0, α the ischemia effect, and *LGE*_*n*_ and *SP*_*n*_ the LGE and SP scores for each sector. It’s known that *Tmax*_*n*_ ≥ 0 and *LGE*_*n*_and *SP*_*n*_ are ∋ [0, 4], then it can be determined *α* ∋ [0, 1]. Since we assumed that the LGE and SP scores were 0 for the healthy volunteer, *C_n_* and *Tmax*_*n*_for the volunteer were equal to *C_H_* and *Tmax_H_*, respectively.

### 2.8 Model optimization

A formal optimization of *C_H_*, *Tmax_H_*, and α were performed where the objective function for the optimization was taken to be the mean-squared errors (MSE) (Guccione et al., 2001). *C_H_* was determined such that the FE model predicted LV end-diastolic volume matched the patient-specific *in vivo* measured volume. *Tmax_H_* was estimated by minimizing the MSE between FE model-predicted and *in vivo* MRI-measured end-systolic longitudinal and circumferential strains and the LV end-systolic volume. The goal of the optimization is to minimize the MSE as follows:

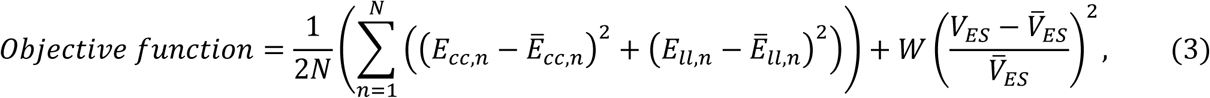

where *n* is the *in vivo* average strain at each sector (note the apex sector was excluded so *n* = 16), *E_cc,n_* the calculated FE circumferential strain, *E_ll,n_* the calculated longitudinal strain, *V_ES_* the LV end-systolic volume and *W* is the weight applied to the volume term. The overbar represents the experimental *in vivo* measurements.

*W* = 10 was applied to the volume term to make the strain and volume effects more balanced. For each case, the *Tmax_H_* was initially = 350 kPa. α was initially = 0.5 in the FMR patient models and = 0 in the healthy volunteer model. No constraints were applied to either *Tmax_H_* or *V_ES_*.

## 3 Results

Patient specific LV and RV pressures are shown in **Table 1** and patient specific LGE, SP and wall motion scores are shown in **Table 2**.

**Table 2.** Patient-specific LGE, SP, and wall motion scores.

|  | Patient 1 |  |  | Patient 2 |  |  |
| --- | --- | --- | --- | --- | --- | --- |
| Segment | LGE score | Stress perfusion score | Wall motion score | LGE score | Stress perfusion score | Wall motion score |
| 1 | 0 | 0 | 0 | 0 | 0 | 0 |
| 2 | 0 | 0 | 0 | 0 | 0 | 0 |
| 3 | 0 | 2 | 1 | 0 | 0 | 0 |
| 4 | 0 | 3 | 3 | 0 | 1 | 0 |
| 5 | 3 | 3 | 4 | 4 | 1 | 2 |
| 6 | 0 | 1 | 1 | 0 | 0 | 0 |
| 7 | 1 | 2 | 3 | 0 | 2 | 0 |
| 8 | 0 | 1 | 0 | 0 | 0 | 0 |
| 9 | 0 | 1 | 0 | 0 | 0 | 0 |
| 10 | 1 | 2 | 3 | 1 | 0 | 1 |
| 11 | 3 | 2 | 3 | 4 | 0 | 3 |
| 12 | 1 | 1 | 1 | 0 | 0 | 1 |
| 13 | 1 | 3 | 3 | 1 | 1 | 0 |
| 14 | 0 | 1 | 0 | 0 | 0 | 0 |
| 15 | 0 | 2 | 3 | 0 | 0 | 1 |
| 16 | 0 | 0 | 1 | 3 | 0 | 3 |
| 17 | 0 | 0 | 3 | 0 | 0 | 0 |

Briefly, Patient 1 was an 88 year old man with multi-vessel CAD (prior left main coronary percutaneous coronary intervention (PCI) with residual chronic occlusion of the left circumflex coronary), heart failure, and severe FMR. CMR demonstrated an enlarged LV with severely reduced LV systolic function (ejection fraction (EF) 30%), subendocardial MI with moderate-severe inducible ischemia involving the inferior and lateral walls as well as subendocardial MI with mild-moderate inducible ischemia involving the LV anterior wall.

Patient 2 was an 84 year old woman with multi-vessel CAD (prior multi-vessel PCI), heart failure, moderate MR. Cardiac MRI demonstrated normal LV size with mildly reduced LV systolic function (EF 45%), transmural inferolateral MI, and mild-moderate inducible ischemia involving the LV anterior wall.

The healthy volunteer had no known heart disease. Note that wall motion was normal and LGE and SP were assumed = 0 for the healthy volunteer.

### 3.1 Mesh convergence study

A mesh convergence study was performed on Patient 1 and Patient 2 to find the minimum number of elements needed to obtain stable calculations of C_*H*_, *Tmax_H_*, and α within the fastest computation time. 4 sets of meshes with different density were created with the transmural elements from 3 to 4, circumferential elements from 40 to 64, and longitudinal elements from 22 to 38. This study determined that 7952 elements (**Table 3**) are required and further mesh refinement only results in an average of 2% change in *C*, < 2% change in *Tmax_H_* and no change in α. Note that only the optimization results for using 7952 elements will be presented in the following sections.

**Table 3.**
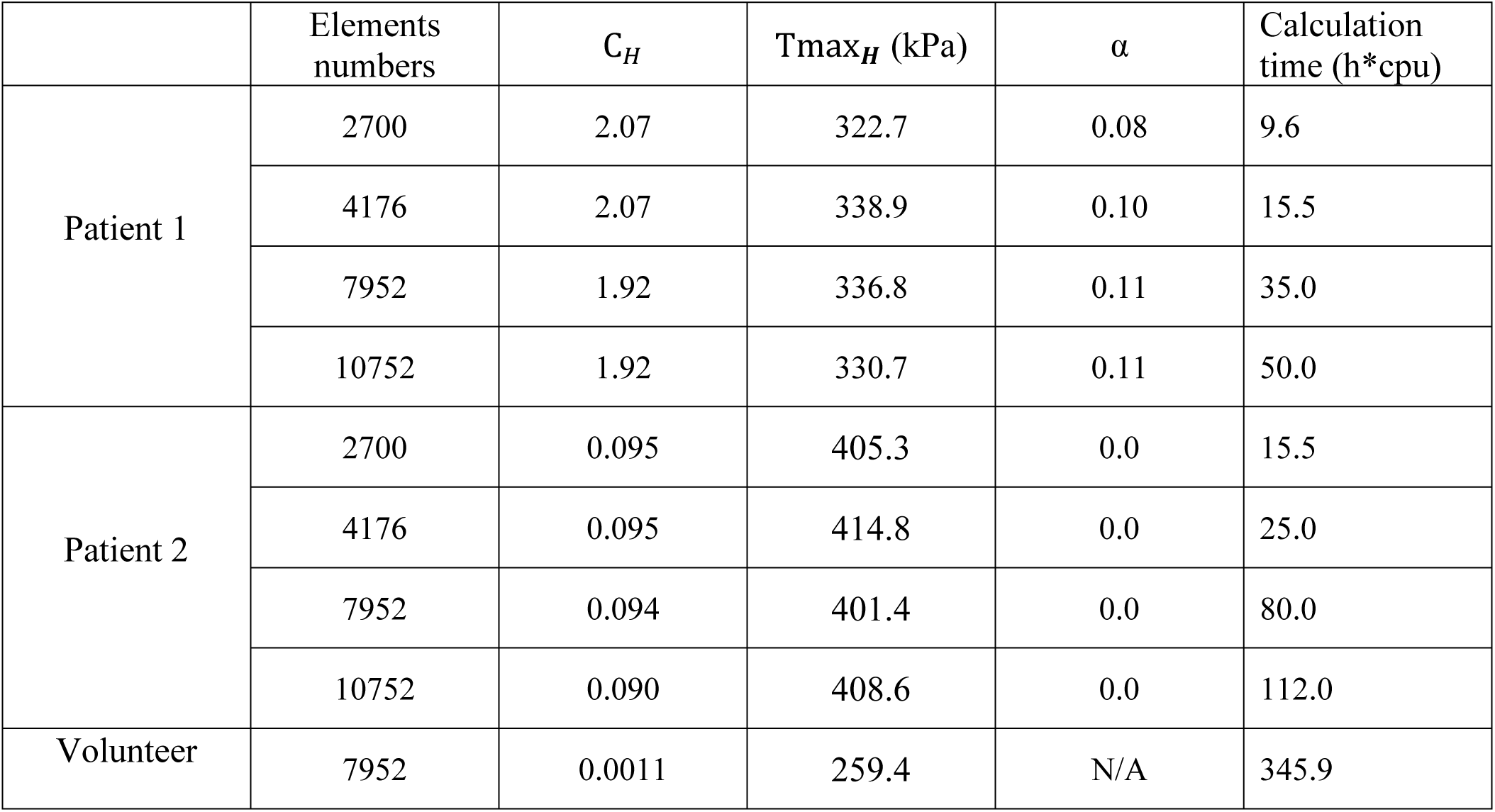
Mesh convergence study.

### 3.2 Testing with synthetic data

Method accuracy was determined using idealized input data. Briefly, two simulations of diastolic inflation and systolic contraction for each patient were firstly conducted by setting α to be 0 and 1, respectively (note that C_*H*_and Tmax_*H*_were set as the optimized values as presented in **Section 3.2**). For each case, the simulated strain and LV ESV were used as input in our optimization framework as “experimental data”. For each case, the optimization was initiated with α initially set at 0.5 and Tmax_*H*_350 kPa. **Figure 4** shows excellent agreement between the synthetically generated “experimental strain” and the strain after optimization. The target and optimized parameters are summarized in **Table 4** with a maximal difference of 1.4%, which shows the proposed method has great capability of predicting Tmax_*H*_ and α accurately.

**Figure 4.**
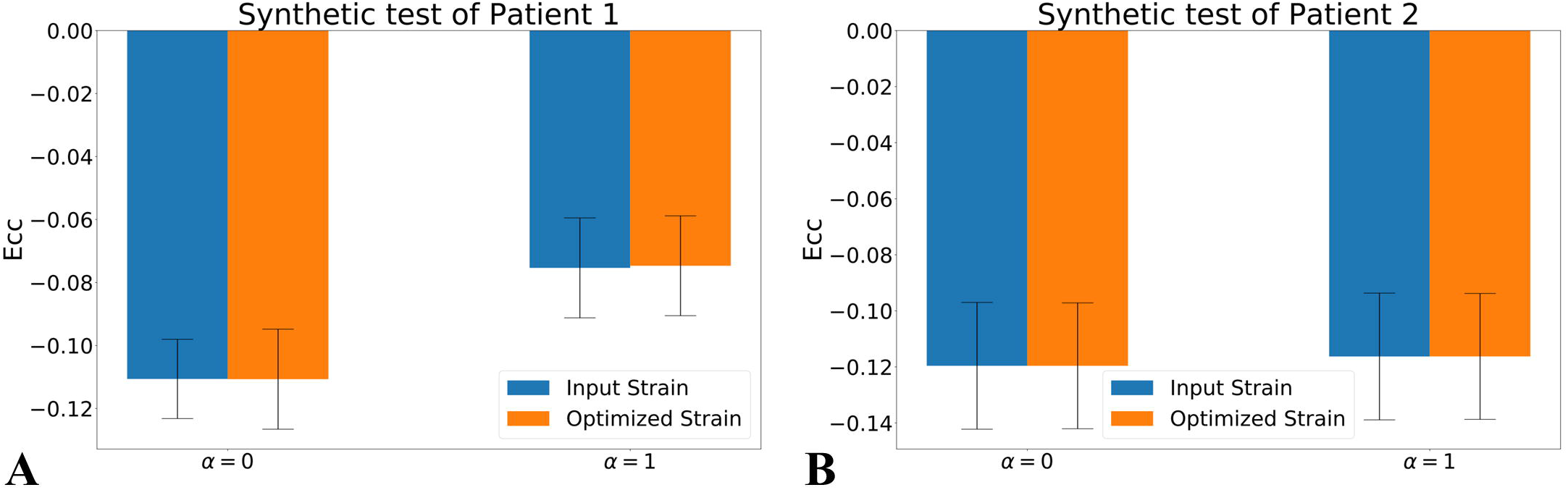
Comparison of the average of *E_cc_* in the synthetic test of Patient 1 (A) and Patient 2 (B).

**Table 4.** The synthetic test.

| ID | Input |  | Optimized |  |
| --- | --- | --- | --- | --- |
| | $Tmax_H$ (kPa) | $\alpha$ | $Tmax_H$ (kPa) | $\alpha$ |
| Patient 1 | 336.8 | 0.0 | 337.6 | 0.0 |
| Patient 1 | 336.8 | 1.0 | 332.2 | 1.0 |
| Patient 2 | 401.4 | 0.0 | 401.5 | 0.0 |
| Patient 2 | 401.4 | 1.0 | 401.0 | 1.0 |

### 3.3 Prediction of C*_H_*, *Tmax_H_*, and α

*C_H_* was determined such that the FE model predicted LV EDV matched the patient-specific *in vivo* measured EDV (**Table 5**).

**Table 5.**
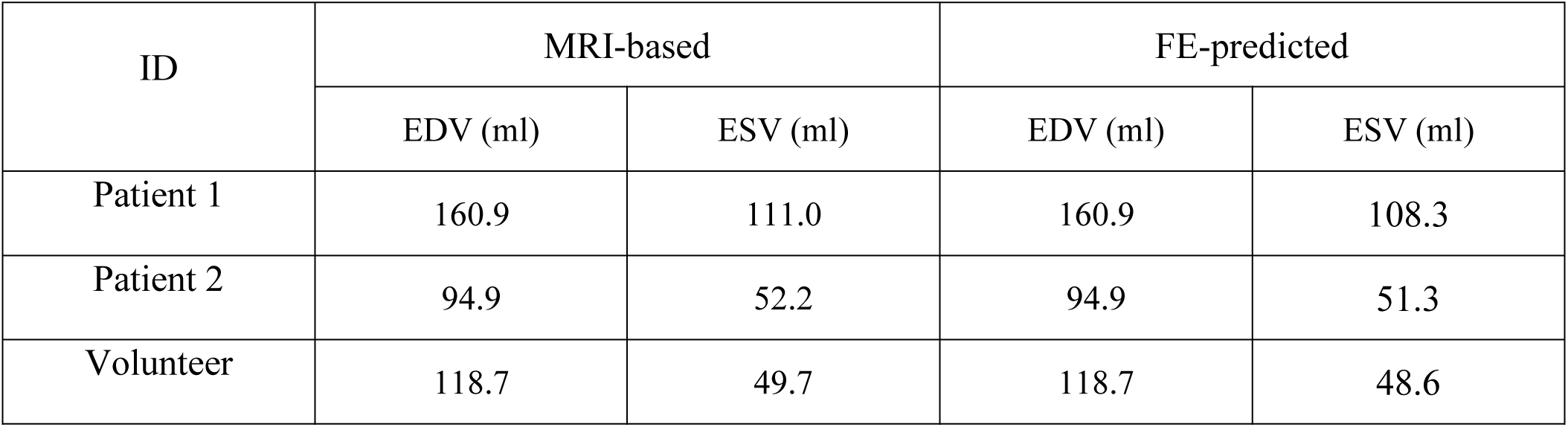
MRI-based *in vivo* volumes and the FE-predicted volumes after optimization.

| ID | MRI-based |  | FE-predicted |  |
| --- | --- | --- | --- | --- |
|  | EDV (ml) | ESV (ml) | EDV (ml) | ESV (ml) |
| Patient 1 | 160.9 | 111.0 | 160.9 | 108.3 |
| Patient 2 | 94.9 | 52.2 | 94.9 | 51.3 |
| Volunteer | 118.7 | 49.7 | 118.7 | 48.6 |

**Figure 5A** shows excellent convergence of the objective function (OF) during a representative model optimization. **Figure 5B** shows a surface plot of MSE of the strain respect to in *Tmax_H_* and α, where the valley of the surface indicates lowest MSE value relating to the best match between the experimental and FE model-predicted strain. The optimized *Tmax_H_* for Patient 1, Patient 2, and the volunteer was 336.8 kPa, 401.4 kPa, and 259.4 kPa and α for Patient 1 and Patient 2 was 0.44 and 0. The optimized *Tmax_H_* and α were obtained with the displaying good convergence as shown in **Figure 6**. The final OF value of 0.45 (an average calculated using all three cases) was obtained indicating generally good agreement between the FE model-predicted systolic strain and the patient-specific *in vivo* measured strain. The LV ESV was accurately predicted at 108.3 ml, 51.3 ml, and 48.6 ml, respectively for Patient 1, Patient 2, and volunteer, which was averagely only 2% smaller than the volume measured from MRI data.

**Figure 5.**
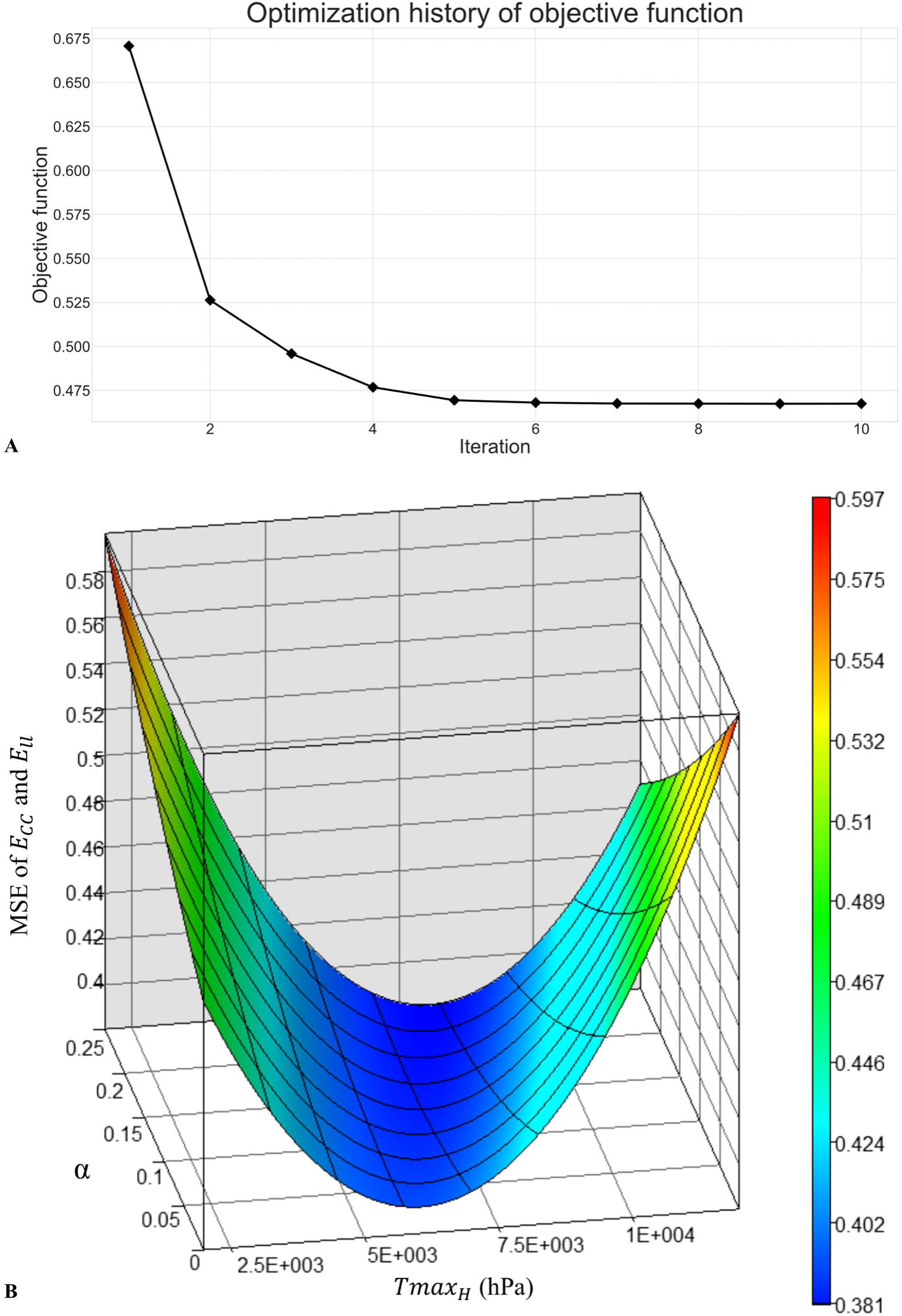
(A) Optimization history of the overall objective function, and (B) surface plot of mean-squared-error of strain respect to *Tmax_H_* and α.

**Figure 6.**
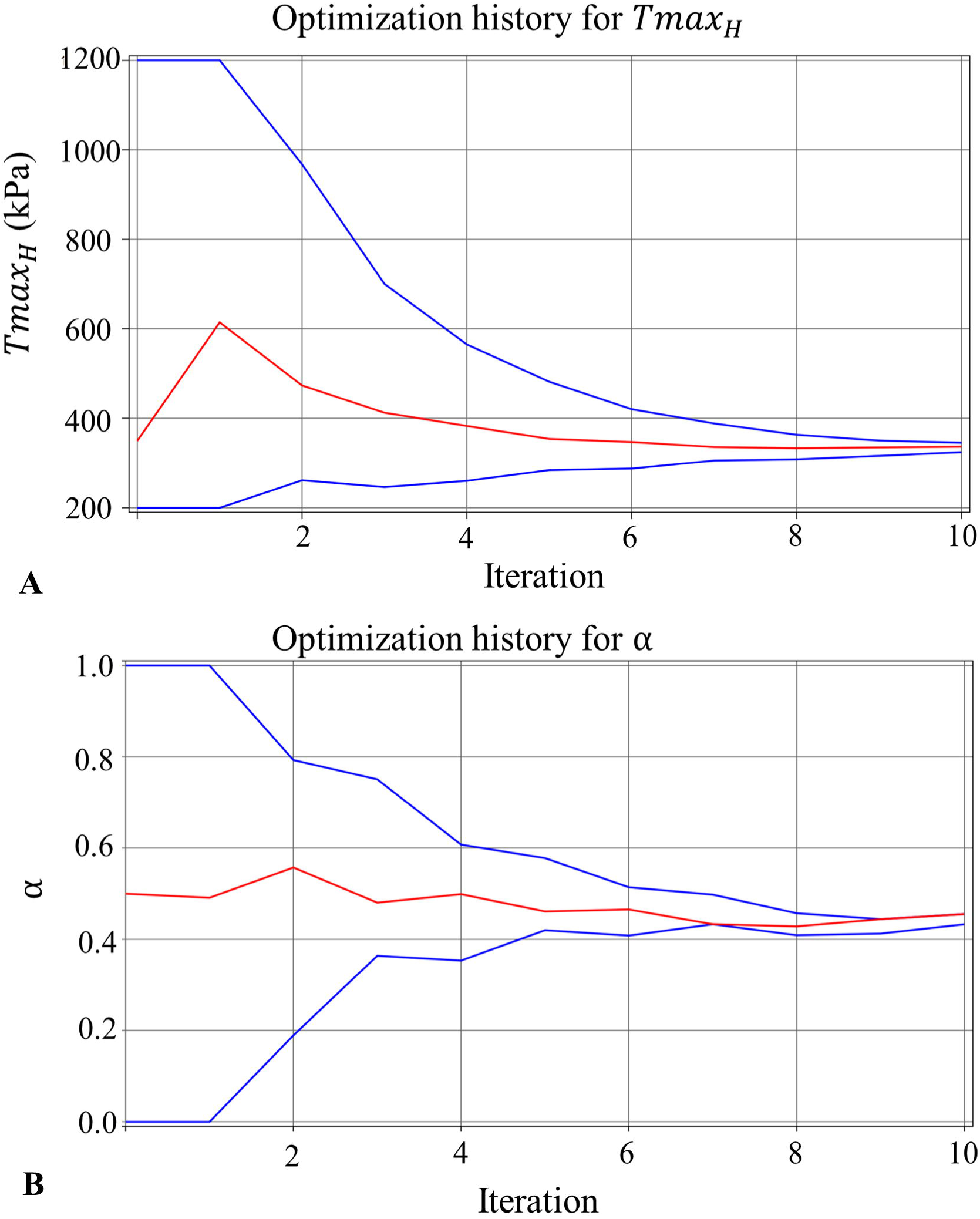
The convergence of the parameter (lines in blue represent the upper and lower bounds) for (A) *Tmax_H_* for healthy myocardium, and (B) ischemia effect, α, on the LV myocardial contractility, with each parameter resulted in a precise final converged optimum.

**Figure 7A** shows a comparison of the average of *E_cc_* across all 16 sectors between the experimental data and calculated with the FE model. In general, FE model predicted strain agreed with experimental data although *E_cc_* was slightly underestimated in Patients 1 2 and slightly overestimated in the volunteer. **Figure 7B** shows that, for Patient 1 and Patient 2, the difference of the average of *E_cc_* between the experimental data and the FE model prediction is significantly reduced when only including the sectors with LGE score < 3.

**Figure 7.**
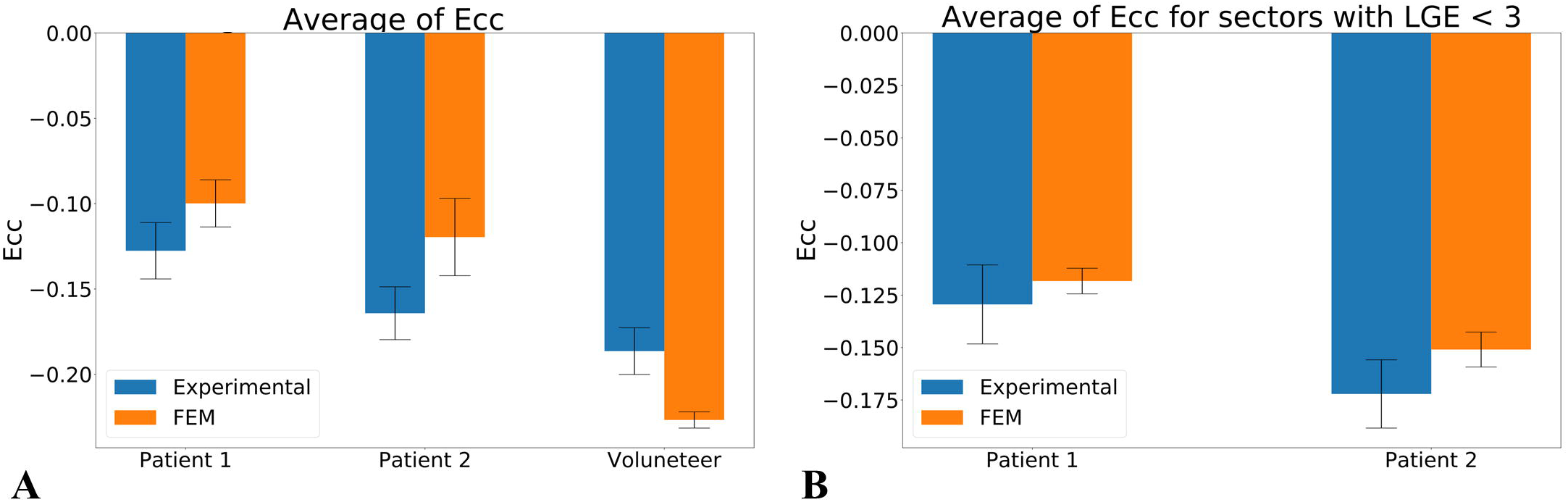
Comparison of the average of *E_cc_* with the standard error of the mean in all three cases for (A) all sectors included and (B) only sectors with LGE < 3.

### 3.4 Virtual REVASC

Given that the FE model predicted a non-zero ischemic effect for Patient 1 (e.g. alpha = 0.44), a virtual REVASC simulation was performed by running a simulation with α reduced to 0 while keeping other parameters unchanged. The comparison of the average of *E_cc_* between baseline and post-REVASC is shown in **Figure 8**, where it can be found that the virtual REVASC improved Patient 1’s *E_cc_* by 10.7%. The LV ESV after REVASC was 102 ml, which was 6% smaller than that at baseline.

**Figure 8.**
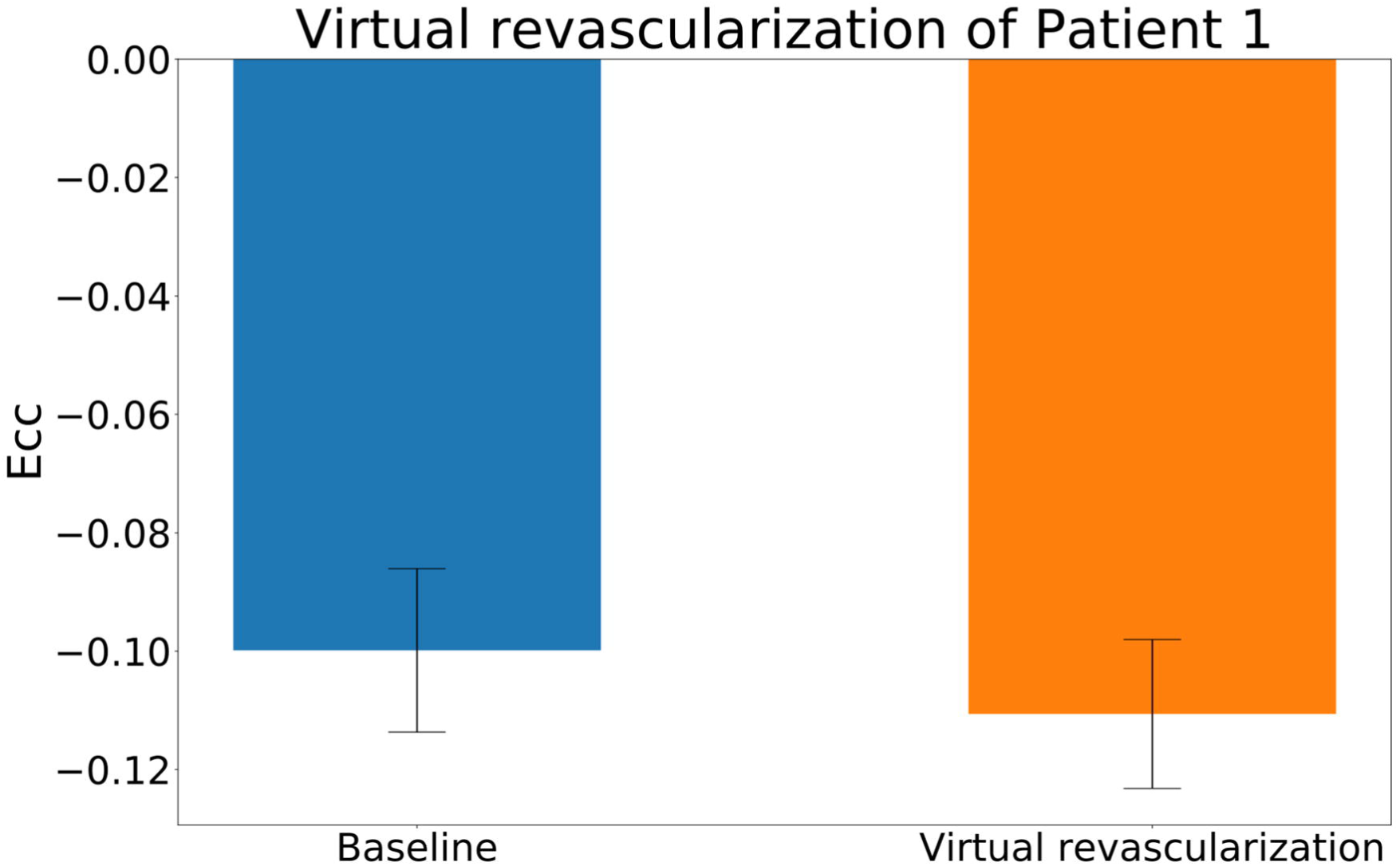
Comparison of the average of *E_cc_* of Patient 1 at baseline and after virtual revascularization.

## 4 Discussion

In this study, we demonstrated a novel method to estimate the effect of ischemia on regional LV myocardial contractility. The proposed method takes the advantage of multiple CMR sequences and the FE modeling to formally optimize the LV mechanics parameters. A linear relationship was proposed to describe the effect of LV infarct and ischemia on LV active contraction. Our approach was tested with CMR data from two patients with multi-vessel coronary disease and moderate FMR and a healthy volunteer. Good agreement between the FE model-predicted systolic strains and the patient-specific *in vivo* measured strains were observed, which suggests that the optimized model was faithful to the experimental data.

### 4.1 Ischemia effect

The effect of ischemia, α, on LV contractility was found to be 0.44 for Patient 1 while 0 for Patient 2. We suggest that this is consistent with the patient specific perfusion data seen in **Table 2** where the cumulative SP score of Patient 1 in sectors with LGE > 3 is 19 while the cumulative ischemia in Patient 2 is only 4.

It should be noted that stress induced perfusion defects are associated with either normal or depressed regional myocardial function and in the latter case the myocardium would be described as hibernating (Wijns et al., 1998). First, by definition, our method identifies only hibernating myocardium and simple demand ischemia without dysfunction is not considered. As a corollary, if a patient with multi-vessel CAD had a large amount of demand ischemia without LV dysfunction, our method would not identify an ischemia effect. Specifically, if the SP defect is showing simple demand ischemia without underlying contractility deficit, then α would likely be 0. If the SP defect is associated with hibernating myocardium where contractility is reduced, then α will likely be > 0.

### 4.2 MI stiffness

Our assumption that infarct stiffness is ten times that of normal myocardium is consistent with our prior studies (Walker et al., 2005; Wenk et al., 2011) and others (Genet et al., 2015), there is reason to believe that better measurement of infarct stiffness is necessary in patients with multi-vessel coronary disease and FMR. First, as seen in **Figure 7B**, model accuracy was much better when sectors with transmural infarction were excluded. Second, the optimized *Tmax_H_* was higher in the two patients with multi-vessel coronary (336.8 and 401.4 kPa respectively) than the healthy volunteer (259.4 kPa). The reason for this discrepancy is unclear but might be explained if **Equation 2** is overestimating the effect of infarct and ischemia on LV contractility. It is anticipated that future studies that measure infarct stiffness in patients with MI but without ischemia will help to better determine the relationship between infarct and myocardial contractility.

### 4.3 Diastolic stiffness in patients with multi-vessel CAD

As seen in **Table 3**, there is a large variance in optimized *C_H_* between the healthy volunteer and patients with multi-vessel CAD and FMR. We suspect this is likely a function of myocardial fibrosis (LGE scores > 0) and, consistent with this, prior clinical studies have shown the extent of LGE to correspond to impaired LV relaxation (Moreo et al., 2009). However, the cumulative LGE scores of Patients 1 and 2 are similar and further studies are therefore necessary to establish our hypothesis. On the other hand, the lack of diastolic strain in this study limits our ability to better optimize passive LV material parameters.

### 4.4 Model accuracy

Results of a mesh sensitivity study suggest that the optimized LV material parameters are relatively insensitive to the mesh density, although slight differences were observed between FE models with different mesh density. **Table 2** indicates that a mesh with 7952 may be the optimal choice for the current study because of the balanced calculation time and parameter optimization accuracy. Furthermore, synthetic test results suggest that the proposed method is capable of precisely determining the LV passive and active material properties and the ischemia effect on LV active contractility.

### 4.5 Virtual revascularization

As listed in **Table 2**, sectors 5 and 11 in Patient 1 had SP score equal to 3. However, expected strain improvement was not found in those sectors after a virtual REVASC. A possible explanation is that sectors 5 and 11 had LGE score equal to 3, which indicated for those sectors, the reduction in contractility was dominated by the LV myocardial infarct. This suggests that for LV myocardium with severe infarction, the benefit from restoring the blood supply is limited.

### 4.6 Limitations

First, only systolic strains were calculated since the CSPAMM data were acquired during the systolic phase, which resulted in a fact that only systolic myocardial material parameters can be optimized.

Because of the lack of diastolic strains, the main diastolic material parameter, *C_H_*, was calibrated to match the measured LV end-diastolic volume. Second, the border zone effect was not taken into consideration in the current study. Third, only a linear relationship between the infarct and ischemia and the LV contractility were employed in this study. Fourth, the LV base boundary conditions may affect the FE-model-based longitudinal strain calculation.

## 5 Conclusion

This study proposed a novel and efficient method to predict the effect of ischemia on the LV contractility with the help of multiple CMR acquisitions and the FE modeling. The proposed method has good agreement between the FE model-predicted systolic strains and the patient-specific *in vivo* measured strains. The proposed method may hold the basis for a better understanding of FMR response to REVASC.

## Acknowledgments

This study was supported by NIH grants R01-HL63348/Ratcliffe and R01-HL128278/Weinsaft. This support is appreciated.

## 7 Author Contributions Statement

YZ, JW and MR designed the study. YZ conducted simulations, analyzed results, and wrote the initial draft of the paper. MR created the finite element models. VW, AM, and JK did data analysis. LG contributed to development of methodology. JW collected the clinical data. LG, JW, and MR supervised the study. YZ, VW, JG, JW and MR contributed to manuscript writing/review.

## 8 Conflict of Interest Statement

The authors declare that the research was conducted in the absence of any commercial or financial relationships that could be construed as a potential conflict of interest.

